# Epigenetic Malleability at Core Promoter Regulates Tobacco PR-1a Expression after Salicylic Acid Treatment

**DOI:** 10.1101/2021.07.24.453639

**Authors:** Niraj Lodhi, Mala Singh, Rakesh Srivastava, Samir V. Sawant, Rakesh Tuli

## Abstract

Tobacco’s *PR-1a* gene is induced by pathogen attack or exogenous application of Salicylic Acid (SA). However, the epigenetic modifications of the most important inducible promoter of the *PR-1a* gene are not understood clearly.
Nucelosome mapping and chromatin immunoprecipitation assay were used to define the histone modification on the *PR-1a* promoter.
Here, we report the epigenetic modifications over core promoter lead to disassembly of nucleosome (spans from −102 to +55 bp,masks TATA and transcription initiation) and repressor complex in induced state. ChIP assays demonstrate repressive chromatin of dimethylation at H3K9 and H4K20 of core promoter maintain uninduced state. While, active chromatin marks di and trimethylation of H3K4, acetylation of H3K9 and H4K16 are increased and lead the induction of *PR-1a* following SA treatment. TSA enhances expression of *PR-1a* by facilitating the histone acetylation, however increased expression of negative regulator (*SNI1*) of *AtPR1*, suppresses its expression in *Arabidopsis thaliana’s* mutants.
Constitutive expression of *AtPR1* in Histone Acetyl Transferases (HATs), LSD1, and SNI1 suggests that its inactive state is indeed maintained by a repressive complex and this strict regulation of pathogenesis related genes is conserved across species.

**SUMMARY:** Histone methylation and acetylation regulation of tobacco *PR-1a* promoter are significant for disassembly of the nucleosome and repressor proteins during induction.

## INTRODUCTION

*PR-1a* (pathogenesis-related-1a) gene is a major defense-related gene of the *PR* family of tobacco (*Nicotiana tabacum*). Linker scanning mutagenesis of the *PR-1a* promoter identified two *as-1* elements and one W-box in the activator region as strong positive, weak negative and strong negative *cis*-elements respectively (Lebel et al., 1998). The core promoter region of *the PR* gene family has a conserved TATA, initiator (INR), and DPE like elements (Lodhi et al., 2008). The detailed chromatin modifications of *PR* gene promoter especially in core promoter sequences during the induction has not been reported yet. Histone modifications dynamically regulate chromatin structure and gene expression for example amino-termini of histones are targets for a series of post-translational modifications including acetylation, methylation, phosphorylation, and ubiquitination (Jenuwein and Allis, 2001; Srivastava and Ahn, 2015; Srivastava et al., 2016; Turner, 2000). Such modifications have been proposed to serve as a ‘histone code’, specifying a chromatin state that determines the transcriptional activity of the genes. Acetylation of histones H3 and H4 is mostly associated with transcriptionally active euchromatin, while methylation is associated with either active or inactive chromatin depending on the methylated amino acid residue (Srivastava et al., 2016; Struhl, 1998), Methylation at H3K4, H3K36, and H3K79 is the hallmark of active transcription, whereas methylation at H3K9, H3K27, and H4K20 is associated with transcriptionally inert heterochromatin (Fischle et al., 2003; Metzger et al., 2005). Lysine can be monomethylated, dimethylated, or trimethylated and each methylation state may have a unique biological function, further increasing the complexity of the ‘histone code’.

In higher plants, dynamic regulation of gene expression by histone methylation and acetylation is still not well understood. One of them, a major study of vernalization in *Arabidopsis thaliana* alters the levels of H3 acetylation and H3K9 and H3K27 methylation in a flowering repressor gene (*FLC*) (Bastow et al., 2004; Sung and Amasino, 2004). Histone acetylation is involved in the regulation of the pea plastocyanin gene (Chua et al., 2003; Sung and Amasino, 2004). Dynamic and reversible changes have also been reported in histone H3K4 methylation and H3 acetylation of rice submergence inducible alcohol dehydrogenase I and pyruvate decarboxylase1 genes in response to the presence or absence of stress, (2006). Unlike reversible histone acetylation process, histone methylation was earlier considered as an irreversible modification. However, Ahmad and Henikoff, (2002) identified a process that removes stable histone methylation through histone exchange. Later, histone demethylases such as LSD1 (Chang and Pikaard, 2005; Metzger et al., 2005) and Jumonji C (JmjC domain-containing) protein (Tsukada et al., 2006; Whetstine et al., 2006; Yamane et al., 2006) were also identified. Four LSD1 like proteins have been reported in *A. thaliana* based on conserved domains (amine oxidase and SWIRM) found on the human LSD1 (Chang and Pikaard, 2005). The LSD1 family is conserved from *S. pombe* to humans and regulates histone methylation by both histone methylases and demethylases. Unlike LSD1, which can only remove mono and dimethyl lysine modifications, JmjC-domain-containing histone demethylases (JHDMs) can remove all three histone lysine-methylation states. (Tsukada et al., 2006), (Yamane et al., 2006),

Nucleosomes at specific positions serve as general repressors of transcription (Lebel et al., 1998; Srivastava et al., 2014). Repressive nucleosomes are remodeled before (Lomvardas and Thanos, 2002) or concurrently (Benhamed et al., 2006) with transcriptional activation. A nucleosome over the TATA region must be displaced to permit the formation of the pre-initiation complex (Lebel et al., 1998; Srivastava et al., 2014). Our present work analyses the modifications in the chromatin architecture of the core promoter region during *PR-1a* gene induction in response to SA. We showed that the modifications in methylation and acetylation states of histones lead to disassembly of the nucleosome and repressor proteins after SA treatment.

## MATERIAL AND METHODS

### Plant materials and growth condition

*Nicotiana tabacum* cv. Petite Havana, used as the wild type, was grown in the greenhouse at 22°C±1 in long-day conditions (16 h light–8 h dark). *Arabidopsis thaliana* Col-0 was used as the wild type. All the mutants were in Col-0 background and *Arabidopsis LSD1* mutants (Chang and Pikaard, 2005) were obtained from the Arabidopsis Biological Resource Center. *Arabidopsis* plants were grown under controlled environmental conditions (19/21°C, 100 μmol photons m^-2^ sec^-1^, 16 h light/8 h dark cycle). Plant accessions used in the study: X12737, X63603, U66264, AT2G14610, AT4G18470, AT5G54420, AT3G47340, ATU27811.

### Antibodies used in ChIP experiment

Antibodies used in ChIP assay were purchased from Santa Cruz Biotechnology: (anti-acetyl histone H4K16, sc-8662, and anti-acetyl histone H3K9/14, sc-8655), Millipore Corporation (anti-monomethyl histone H3K4 (07-436), anti-dimethyl histone H3K4 (07-030), anti-trimethyl histone H3K4 (07-473), anti-monomethyl histone H3K9 (07-450), anti-dimethyl histone H3K9 (07-441), anti-trimethyl histone H3K9 (07-442), anti-monomethyl histone H4K20(07-440), anti-dimethyl histone H4K20 (07-367), anti-trimethyl histone H4K20 (07-463), and anti-histone H3 (06-755), CoREST, HDAC1 and Arabidopsis anti-LSD1 (developed in our lab).

### Plasmid constructions and plant transformation

The *PR-1a* promoter was amplified from the genomic DNA of tobacco by using forward PRF and reverse PRR primers (**Table S1**) and fused to *gusA* gene in pBluescript SK^+^ as in Lodhi *et al*, (2008) (Lodhi et al., 2008). *Agrobacterium tumefaciens* mediated plant transformation was performed comprising construct containing *PR-1a* promoter to examine the expression in stable transgenic lines of *Nicotiana tabacum* cv. Petit Havana.

### SA and TSA treatments of plant leaves

The effect of salicylic acid (SA) (Sigma, USA) and Trichostatin A (TSA) (Sigma, USA) on promoter expression were studied on discs. Discs of 3 cm diameter were excised from expanded leaves of transgenic plants and floated on water or 2 mM SA in petri-dish. For inhibition of histone deacetylase, the leaves were treated with 300 μM TSA. The leaves were incubated for 12 h in light at 25± 2 °C. In the case of *A. thaliana*, 100 mg intact 21-days old plantlets were floated on water or SA.

### Determination of GUS enzymatic activity

The leaf discs were ground in liquid-nitrogen and extracted with 1 ml GUS assay buffer (50 mM Na_2_HPO_4_ pH 7, 10 mM EDTA, 0.1% (v/v) Triton X-100, 1 mM DTT, 0.1% (w/v) N-lauryl sarcosine and 25 μg/ml phenylmethylsulfonyl fluoride. The extract was centrifuged at 16,000 x g for 20 min at 4°C. After centrifugation, 90 μl supernatant was mixed with 10 μl GUS assay buffer containing 1 mM of 4-methylumbelliferyl-β-D-glucuronide (MUG) as substrate. The mixture was incubated at 37 °C for 1 h. The product 4-methylumbelliferon (MU) was quantified using a fluorimeter (Perkin Elmer LS55, USA). Protein concentration was determined by Bradford assay (Bio-Rad, Hercules, CA, USA) GUS activity is expressed in units (1 unit = 1 nmol of 4-MU/h/mg of protein).

### DNA Sequence mapping of nucleosome’s border

The 10 g leaves were treated with water or 2mM SA for 12 h with gentle agitation in light. After 12 h, the samples were subjected to cross-linking in NIB1 buffer (0.5 M hexylene glycol, 20 mM KCl, 20 mM PIPES at pH 6.5, 0.5 mM EDTA, 0.1% Triton X-100, 7 mM 2-mercaptoethanol) in the presence of 1% formaldehyde for 10 min. The cross-linking was stopped by adding glycine to a final concentration of 0.125 M for 5 min at room temperature. The leaves were then rinsed with water, ground to powder in liquid nitrogen, and treated with nuclei isolation buffer NIB1. The extract was filtered through 4 layered muslin cloth and finally filtered sequentially through 80, 60, 40 and 20 μm mesh sieves. The filtrate was centrifuged at 3,000 x g at 4 °C for 10 min. The pellet was suspended in NIB2 (NIB1 without Triton X-100) and centrifuged again. The pellet was suspended in 5% percoll, loaded on 20-80% percoll (U.S. Biologicals, USA) step gradient and centrifuged. The nuclei were removed from the 20-80% percoll interface, washed in NIB2, and resuspended in NIB1 buffer. The nuclear preparation equivalent to A_260_ of 100 was incubated with micrococcal nuclease (300 units/μl) (Fermentas, USA) in a buffer containing 25 mM KCl, 4 mM MgCl_2_, 1 mM CaCl_2_, 50 mM Tris-Cl at pH 7.4 and 12.5% glycerol at 37 °C for 10 min. The reaction was stopped by adding an equal volume of 2% SDS, 0.2 M NaCl, 10 mM EDTA, 10 mM EGTA, 50 mM Tris-Cl at pH 8 and treated with proteinase K (100 μg/ml) (Ambion, USA) for 1 h at 55 °C. The crosslink was reversed by heating at 65°C overnight. The DNA was extracted by phenol: chloroform and precipitated in ethanol. The DNA was separated on 1.5 % agarose gel and fragments of an average size of 150 bp were purified, denatured, and hybridized with 20 ng of end labeled forward PF3 and reverse NR1 primers of region 1. Primer extension was performed at 37°C using 13 units of sequenase (U.S. Biologicals, USA) in 1x sequenase buffer containing 0.01 M DTT and 0.1 mM dNTPs according to manufacturer’s protocol including ladders of all four nucleotides. The products were analyzed in 8% sequencing gels. The sequences of primers used in primer extension are given in **Table S1**.

### Detection of nucleosomes on tobacco *PR-1a* promoter using ChIP DNA template

Standard PCR was used to locate nucleosomes in the upstream, downstream, and core promoter regions. MNase digested mononucleosome DNA precipitated with H3 was used as a template to detect the amplicon in uninduced, induced state and TSA treated leaves. Mononucleosomes were purified using Hydroxyapetite (HAP) protocol (Brand et al., 2008). The forward primers (PF3, NPAF1, NPAF5, and NPCF1) and the reverse primers (NR1, NPAR1, NPAR5, and NPCR1) were used to analyze the protection of the core promoter (−102 to +55 bp), the upstream (−362 to −213 and −262 to −102 bp) and downstream (+59 to +208 bp) regions respectively of the *PR-1a* promoter against micrococcal nuclease digestion in the uninduced and induced states. The sequences of all primers are given in **Table S1**. To do native ChIP, 1.5 – 2 g leaf discs of tobacco excised from 8-9 week old plants were floated on water or 2mM SA and 300 μM TSA for 12 h with gentle agitation in the light. After 12 h the samples were rinsed with water and ground into powder in liquid nitrogen. Nuclei were extracted and washed with 1 ml of buffer N (15 mM Trizma base, 15mM NaCl, 60mM KCl, 250mM sucrose, 5mM MgCl_2_, 1 mM CaCl_2_,pH-7.5, 7 mM 2-mercaptoethanol, 1 mM phenylmethylsulfonyl fluoride, and 50 μl/ml plant protease inhibitor cocktail) (Sigma chemicals, USA). Thereafter nuclei were suspended in 100μL buffer N. DNA content was estimated in a 10μl aliquot and MNase treatment were given using 1unit/μg DNA for 10 min at 37°C., and finally eluted in 300μL of HAP elution buffer (500mM NA_2_PO_4_ pH7.2, 100mMNaCl, 1mM EDTA) and was diluted with 1700μL of ChIP dilution buffer (1.1% Triton X-100, 1.2 mM EDTA, 16.7 mM Tris-HCl, pH 8, 167 mM NaCl, and 50 μl/ml protease inhibitor cocktail). The diluted chromatin solution was then subjected to 1 h of pre-cleaning treatment at 4°C with 80 μl of salmon sperm DNA/protein agarose (Upstate; 16-157). An aliquot of 50 μl was removed for the total input DNA control. Immunoprecipitation was performed overnight (18 h) at 4°C using 600 μL chromatin solution with histone H3 antibodies (typically at 1:150 final dilutions) or without antibodies (mock control). Immunoprecipitates were collected after incubation with 40 μL of salmon sperm DNA/protein agarose (50% suspension in dilution buffer) at 4°C for 1 h. The protein A agarose beads bearing immunoprecipitate were then subjected to sequential washes and eluted twice with 250 μL elution buffer each time (1% SDS and 0.1M NaHCO_3_). For the input DNA control (50 μL), 450 μL elution buffer was added. Protein was removed by 1.1 μL proteinase K (20mg/ml) at 45° C for 1h and RNA by 2μL of RNaseA (1mg/ml) digestion at 37°C for 1h. The DNA was purified by phenol: chloroform extraction and ethanol precipitation. Purified DNA was resuspended in 50 μL TE buffer for PCR analysis.

### Southern hybridization to detect nucleosomes in promoter region of tobacco *PR-1a*

Twenty micrograms of purified MNase-digested DNA was analyzed to find out the position of nucleosomes in *PR-1a* promoter. Eight probes of 200 bp from the core promoter region were designed (R1 to R8). For positive control, 10 pg *PR-1a* promoter (PCR amplified) and for negative control 10 pg of sonicated calf thymus DNA was used. The entire DNA was transferred on to nylon membrane and incubated at 42°C overnight with a probe.

### ChIP PCR using precipitated DNA with different antibodies

The leaf discs (1.5 – 2 g) of tobacco excised from 8-9-week-old plants were floated on water or 2mM SA for 12 h with gentle agitation in light. After 12 h the samples were subjected to 1% formaldehyde cross-linking in a cross-link buffer (0.4 M sucrose, 10 mM Tris-HCl, pH 8, and 1 mM EDTA) under vacuum for 10 min. Formaldehyde cross-linking was stopped by adding glycine to a final concentration of 0.125 M and incubating for 5 min at room temperature. The leaf pieces were then rinsed with water and ground to powder in liquid nitrogen. Nuclei were extracted and lysed with 300 μl of lysis buffer (50 mM Tris-HCl, pH 8, 10 mM EDTA, 1% SDS, 1 mM phenylmethylsulfonyl fluoride, 10 mM Sodium butyrate, 1 mM benzamidine, and 50 μL/ml protease inhibitor cocktail) (Sigma chemicals, USA). The resulting chromatin was subjected to pulse sonication (six pulses, 95% power output for eight times) using a Bransonic M3210 (Danbury, USA) to obtain DNA fragments with size ranging from 500 to 1000 bp. After sonication, a 25 μl aliquot was removed for the total input DNA control, and the rest of the chromatin solution was diluted 10 times with ChIP dilution buffer (1.1% Triton X-100, 1.2 mM EDTA, 16.7 mM Tris-HCl, pH 8, 167 mM NaCl, and 50 μl/ml protease inhibitor cocktail). The diluted chromatin solution was then subjected to 1 h of precleaning treatment at 4°C with 40 μl of salmon sperm DNA/protein agarose (Upstate; 16-157) (50% suspension in dilution buffer without Na butyrate and protease inhibitor cocktail) to reduce nonspecific interactions between protein-DNA complexes and the agarose beads. Immunoprecipitation was performed overnight (18 h) at 4°C using 600 μL chromatin solution with antibodies (typically at 1:150 final dilutions) or without antibodies (mock control). Immunoprecipitates were collected after incubation with 40 μL of salmon sperm DNA/protein agarose (50% suspension in dilution buffer) at 4°C for 1 h. The protein A agarose beads bearing immunoprecipitate were then subjected to sequential washes and eluted twice with 250 μL elution buffer (1% SDS and 0.1M NaHCO_3_). Samples were then reverse cross-linked at 65°C under high salt (0.2 M NaCl) for 6 h. For the input DNA control (25 μL), 275 μL TE buffer (10 mM Tris-HCl, pH 8, and 1 mM EDTA) was added and reverse cross-linked. After reversing the cross-links, the protein was removed by 1.1 μL of proteinase K (20mg/ml) at 45° C for 1h and RNA by 2μL of RNaseA (1mg/ml) (Qiagen) digestion at 37°C for 1h. The DNA was purified by phenol: chloroform extraction and ethanol precipitation. Purified DNA was resuspended in 40 μL TE buffer for PCR analysis.

For ChIP PCR the target region of the *PR-1a* promoter was −102 to +55 with reference to the transcription start site. Forward primer PF3 and reverse primer NR1 were used for amplifying the core promoter. Tobacco *ACTIN* promoter was taken as an internal control for active chromatin, using forward AGF and reverse AGR primers for PCR **Table S1.** For testing the enrichment of various modifications on the R8 promoter at different time points, forward primer PF3 and reverse primer NR1 were used for qRT-PCR. Reactions were placed in 25 μl volume in triplicate according to the manufacturer’s instruction (Invitrogen SYBR green ER) on ABI PRISM 7500.

### Transcripts detection of different defense related genes of *Arabidopsis* using Quantitative real-time PCR

To compare the transcript levels of *AtPR1*, *AtSNI1*, *AtPDF1.2*, and *AtASN1*, leaves of wild type *A. thaliana* plants (Col-0) were treated with water, 2 mM SA, TSA alone or TSA and SA both. After 12 h, total RNA isolated by Tri-reagent (Sigma) and treated with RNase free DNase. For temporal expression of tobacco *PR-1a*, the total RNA was isolated from the leaf discs which were floated on SA for different time periods. The first strand cDNA was synthesized, using 2 μg RNA, as per manufacturer’s instructions (Invitrogen, USA). Quantitative real-time PCR (qRT-PCR) was used to determine the expression of *AtPR1* in uninduced and induced states, using forward ATPRF and reverse ATPRR primers. Tobacco *PR-1a* expression at different time points was followed by standard PCR by using forward NPRF and reverse NPRR primers. The *AtACTIN7* and *UBIQUITIN* genes were used as an internal control. The sequences of primers used are given in **Table S1**.

## RESULTS

### Nucleosome over core promoter of *PR-1a* spans from −102 to +55 bp in uninduced state

Earlier, we reported a distinct nucleosome over the core promoter region of *PR-1a* in the uninduced state disassembles upon SA induction to initiate the transcription (Lodhi et al., 2008). In the present study, we reported mapping of nucleosome using a primer extension method. It was performed after confirming the presence of nucleosome as well as on entire promoter of *PR-1a* by southern hybridization. We performed by deviding the entire length of the *PR-1a* promoter (1.5Kb) into eight distinct regions of around 200 bp (R1 to R8). The region encompassing the core promoter and transcription start site (TSS) was designated as R8 (Fig. S1). The mono-nucleosome template from uninduced tobacco plants was prepared by digesting with micrococcal nuclease (MNase) enzyme (digest the linkler region). Probes from different regions of the *PR-1a* promoter were used in southern hybridization with the MNase digested mono-nucleosome template. Southern hybridization reveals the presence of nucleosomes over five regions including R1, R2, R4, R5 and R8 on promoter (Fig. S1). Nucleosome boundaries of R8 nucleosome (over core promoter) were mapped using the primer extension method with forward (PF3) and reverse (NR1) primers (Fig. 1A is showing the sequencing with one primer). The boundaries of the nucleosome were found to be spanning from −102 to +55 bp (with respect to the TSS) in *PR-1a* (Fig. 1B). The nucleosome over R8 masked the TATA region, transcription initiation site (+1), and downstream promoter region (−102 to +55 bp) in the uninduced state of *PR-1a*.

**Fig. 1.**
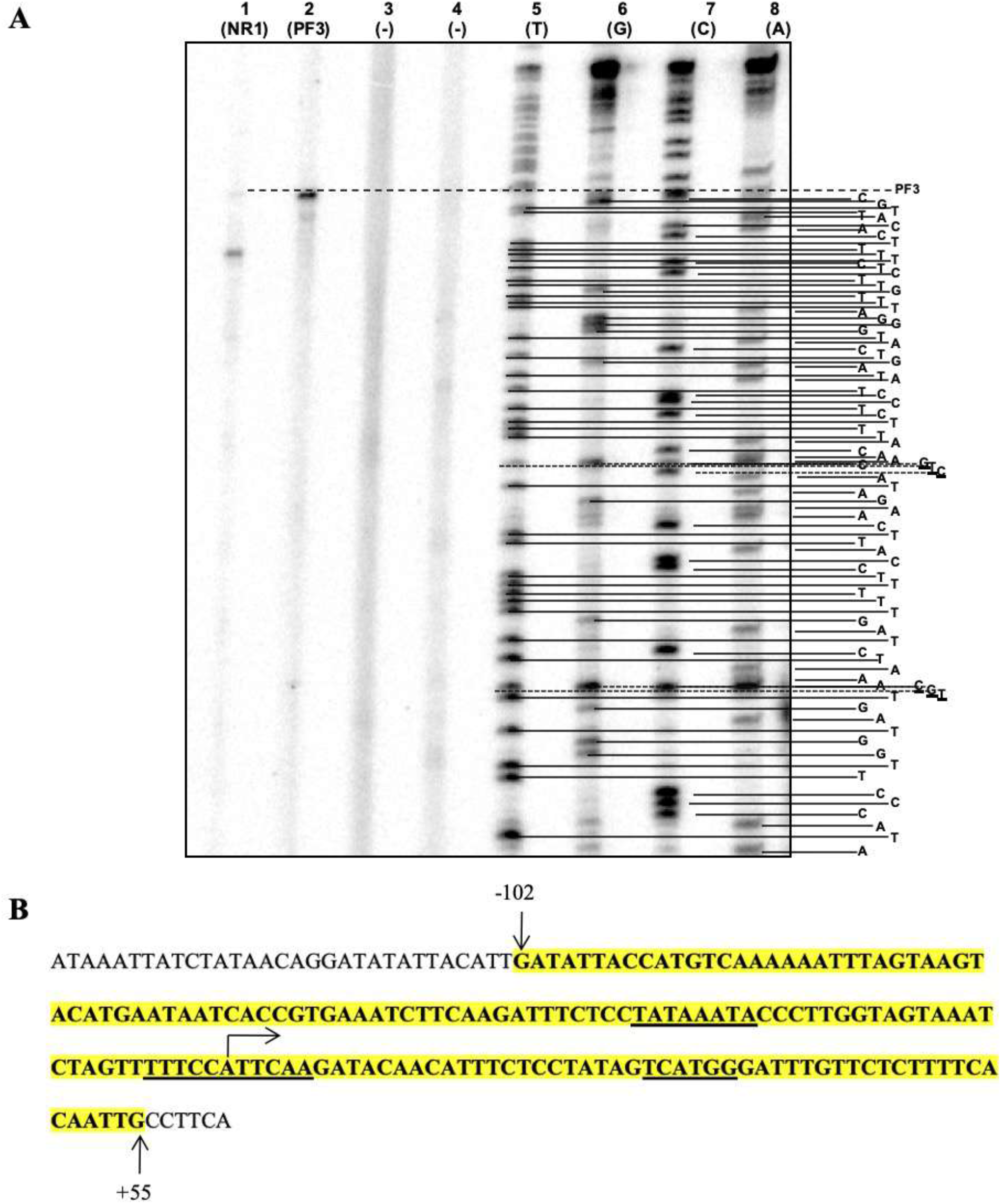
Mapping border sequences of the core nucleosome. (A) Lane 1: Amplified product of reverse primer (NR1). Lane 2: Amplified product of forward primer (PF3). Lane 3 and 4: non template controls for NR1 and PF3 (negative controls). Lanes 5 to 8: sequence ladders for T, G, C and A respectively. (B) Nucleotide sequence of core promoter of tobacco *PR-1a* promoter showing in bold the −102 to +55 region covered by the nucleosome. The TATA, *Inr* like region and downstream promoter like sequences are underlined and TSS is showed by arrow.

### Histone acetylation (H3K9Ac and H4K16Ac) marks associated with the activation of *PR-1a* followed by SA treatment

It was demonstrated that the *PR-1a* induction coincided with the disappearing or disassembly of the nucleosome over region 8 (R8) (Lodhi et al., 2008). Here, we further examined the epigenetic changes in chromatin responsible for the disassembly of the nucleosome. Since histone acetylation associated with transcriptional activation and histone acetylation of H3K9/14 and H4K16 has been demonstrated in the activation of genes (Santos-Rosa and Caldas, 2005; Shahbazian and Grunstein, 2007; Shogren-Knaak and Peterson, 2006). We checked the acetylation status of the H3K9 and H4K16 of R8 nucleosome using the ChIP approach in a time-dependent manner. ChIP results showed that the onset of transcription of *PR-1a* strongly correlated with the H3K9/14 and H4K16 acetylation. Acetylation of these lysine residues increased gradually from 3 to 9h post-SA treatment reaching maximum at 9h (Fig. 2A and B),.

**Fig. 2.**
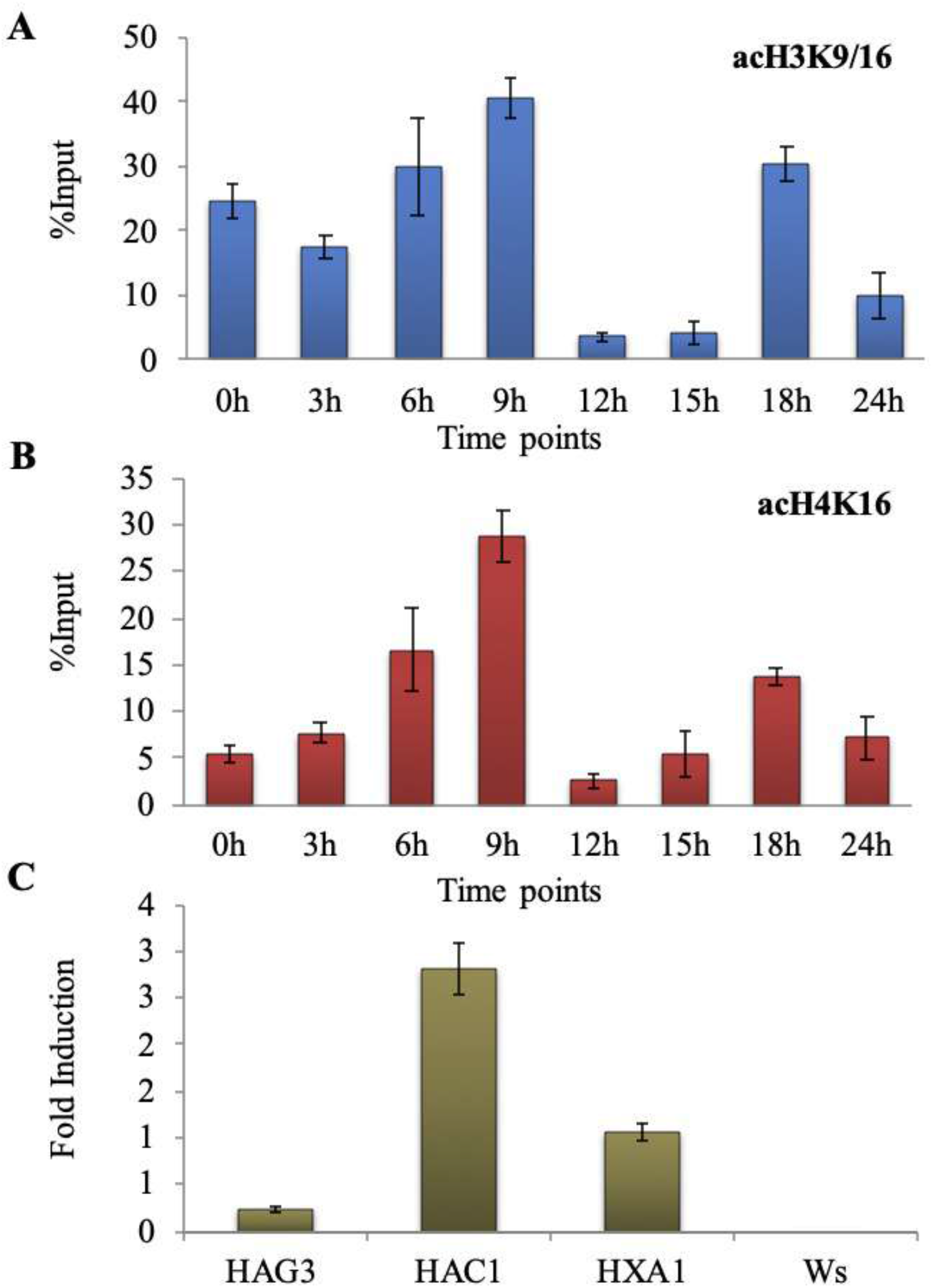
Time course analysis of the acetylation chromatin state on the tobacco *PR-1a* core promoter region in uninduced and induced state, and *AtPR1* expression in *Arabidopsis* HAT mutants. (A) Histone acetylation status on R8 was analyzed by ChIP assay using A) anti-acetyl H3K9, B) anti-acetyl H4K16 antibodies. The immunoprecipitated DNA was analyzed by qRT-PCR. The histogram represents the % input (Y-axis) at different time points (X-axis) with SD. Constitutive expression of *AtPR1* in HAG3, HAC1, HXA1 gene mutant lines of *Arabidopsis* in comparison to wild type (Ws) in uninduced state and quantitative (C)*. AtPR1* expression was quantified by real-time PCR.

To examine whether *PR1* locus genetically interacted with histone acetyltransferases, we performed experiments in *Arabidopsis thaliana* (Ws) because histone acetyltransferase mutants’ plants of tobacco were not available and also assuming histone acetyltransferases are conserved across the plant species. Three HAT mutants of *Arabidopsis thaliana* (Ws) i.e. HAG3, HAC1, and HXA1 were examined. The estimation of *AtPR1* transcript in the mutants in uninduced states showed a significant increase of transcript as compared to the wild type. An increase in the uninduced expression of *AtPR1* suggested the loss of stringent regulation of *PR1* in the uninduced condition (Fig. 2C). Therefore, histone acetyltransferases HAG3, HAC1, and HXA1 are essential to acetylate histone marks of nucleosome to activate the transcription of *AtPR1*.

### Nucleosome disassembly of *PR-1a* core promoter is required for transcriptional activation

The disappearance of the nucleosome could be either due to nucleosome sliding or complete disassembly. To further understand the fate of nucleosome remodeling at the PR-1a core promoter, we addressed the histone H3 occupancy either on Group 3 (R8) or on flanking upstream Group 1 or 2 and downstream promoter region group 4 of *PR-1a* by ChIP using anti-H3 antibody in uninduced and SA treated leaves. We observed distinct nucleosome over group 3 (R8, −102 to +55) as evident PCR amplified product in case of uninduced control (Fig. 1). Since SA treatment affects histone acetylation of the nucleosome over *PR-1a* core promoter region upon the induction (Fig. 2), the histone H3 occupancy was also studied in Trichostatin A (TSA, an inhibitor of histone deacetylase) treated tobacco leaf discs in the presence or absence of SA (Fig. 3). The nucleosome over group 3 disappeared with SA induction, however treatment with TSA in the presence or absence of SA inhibited nucleosome disappearence (Fig. 3A). We did not observe nucleosome protection over group 2 (−213 to −102) in any of the conditions tested indicating the lack of nucleosome over this region (Fig. 3B).

**Fig. 3.**
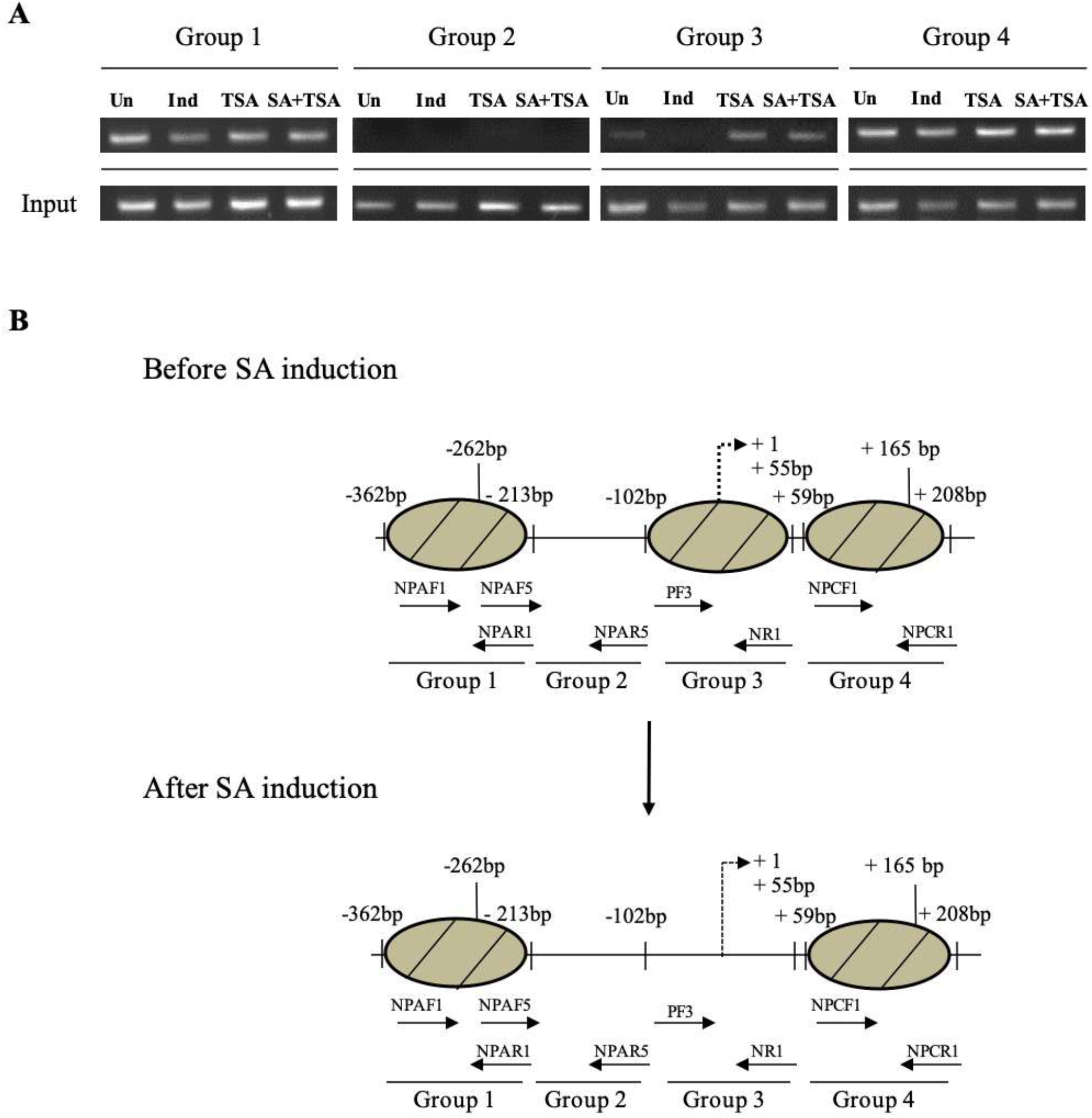
Nucleosome mapping on the *PR-1a* core promoter region in uninduced, SA, TSA and SA + TSA treated leaves by anti-H3 ChIP-PCR. (A) Standard PCR was done to detect nucleosomes in the core promoter (−102 to +55 bp) and flanking upstream (−102 to −213 bp and −213 to −362 bp) and downstream (+59 to +319 bp) regions of the native *PR-1a* promoter. The Input DNA is used as ChIP control (for each primer set) as shown below each lane. (B) The models depict the location of nucleosomes on *PR-1a* promoter before and after SA induction upon the regions analyzed in (A).

Multiple sets of primer pairs and ChIP template DNA were used to detect nucleosomes associated with different regions of the core promoter and flanking promoter of *PR-1a* (Fig. 4). Therefore, results suggest a explanation for the repression of *AtPR1* transcript with TSA treatment in *A. thaliana* (discussed later). The promoter flanking region in group 1 (−362 to −213) and group 4 (+59 to +208) also have distinct nucleosomes, however, these nucleosomes did not show any change post-SA or TSA treatment.

**Fig. 4.**
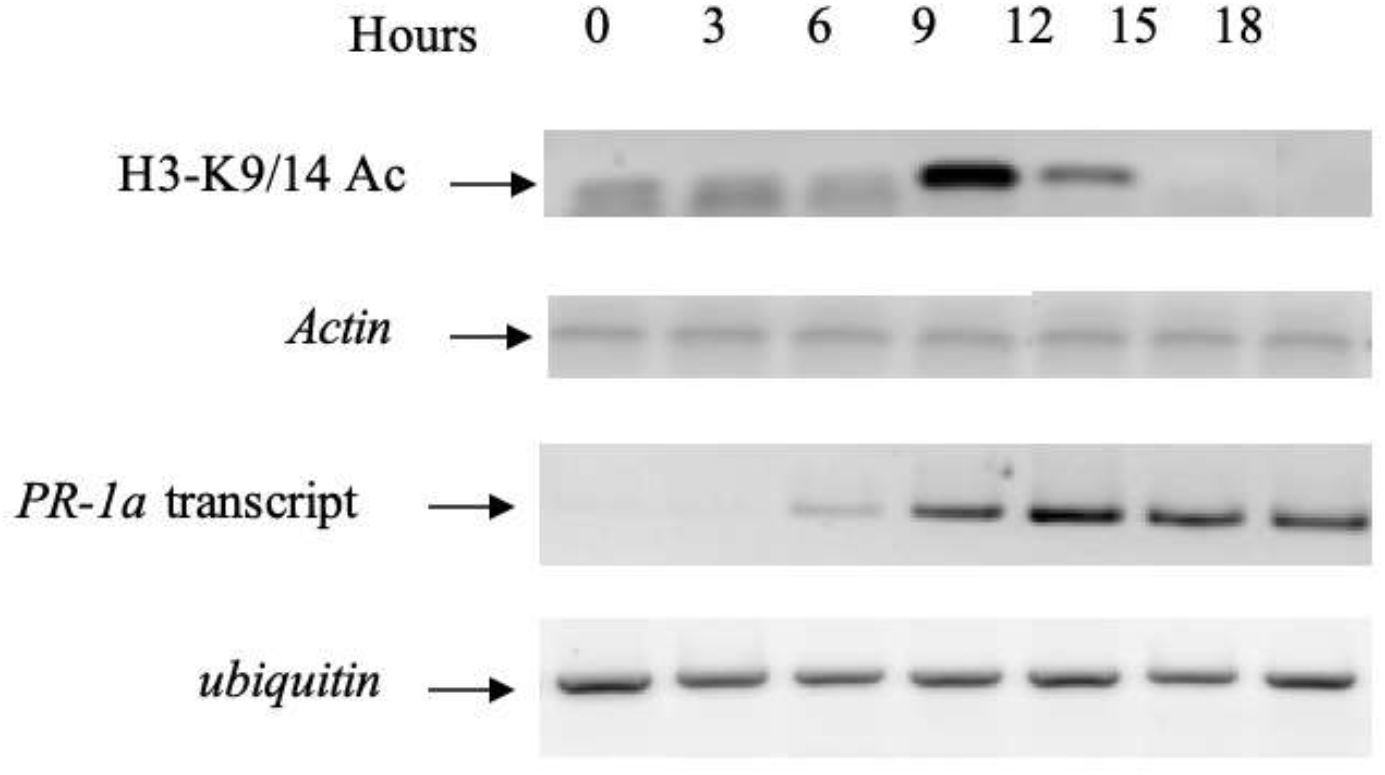
Temporal changes in histone acetylation at *PR-1a* core promoter in relation to *PR-1a* transcript. The ChIP assay was performed on tobacco leaves floated on SA for 3 to 18 h, using antibodies against acetylated H3K9/14. The PCR products from immunoprecipitated DNA (correspond to core promoter region) are shown at different time points. The input template was used as control. Tobacco *PR-1a* transcripts were estimated by RT-PCR at different time points, following SA induction. Tobacco *UBQ* was used as an internal control for transcript analysis.

### Temporal regulation of *PR-1a* by SA treatment correlates with the acetylation of H3-K9/14 of core promoter nucleosome

The *PR-1a* is a late inducible promoter, its activation was noticed 9 h after SA treatment (Fig. 4). To further understand the correlation between *PR-1a* transcription and acetylation of its promoter, we studied the temporal regulation of *PR-1a* in response to SA and performed ChIP with acetylated H3K9/14 and tri-methylated H3K4 antibodies on the core promoter at different time points. The onset of transcription of *PR-1a* was correlated strongly with the acetylation of H3K9/14 (Fig. 4). The H3K9/14 was highly acetylated at 9 h, remained so till 12 h post-SA treatment, and then declined. The results indicated that during 9 to 12 h post-SA treatment, there was a sharp, though the transient increase in acetylation of H3 in the nucleosome of the core promoter, whereas a slight increase in tri-methylation of H3K4 from 6 to 9 h post-SA treatment.

### TSA enhances early expression of tobacco *PR-1a* followed by SA treatment

The effect of HDAC inhibitor TSA was examined on the expression of *PR-1a*. The leaves were treated with SA for 4h (to get the induction signal) and then shifted to either water or TSA. The expression of *PR-1a* promoter was examined by assaying the *GUS* reporter gene fused to it. The analysis of three independent transgenic lines showed a clear effect of TSA. The expression was higher in the TSA treated leaves till 25 h in comparison to the water treated leaves (Fig. 5). Higher expression correlated well with the H3K9/14 acetylation (Fig. 4). After 25 h, there was no difference in expression in the two cases. The results indicated that short exposure to SA leads to transcription of *PR-1a* which was vulnerable to suppression by HDACs. However, after 25 h in water or TSA, stable H3K9/14 acetylation-insensitive expression was noticed.

**Fig. 5.**
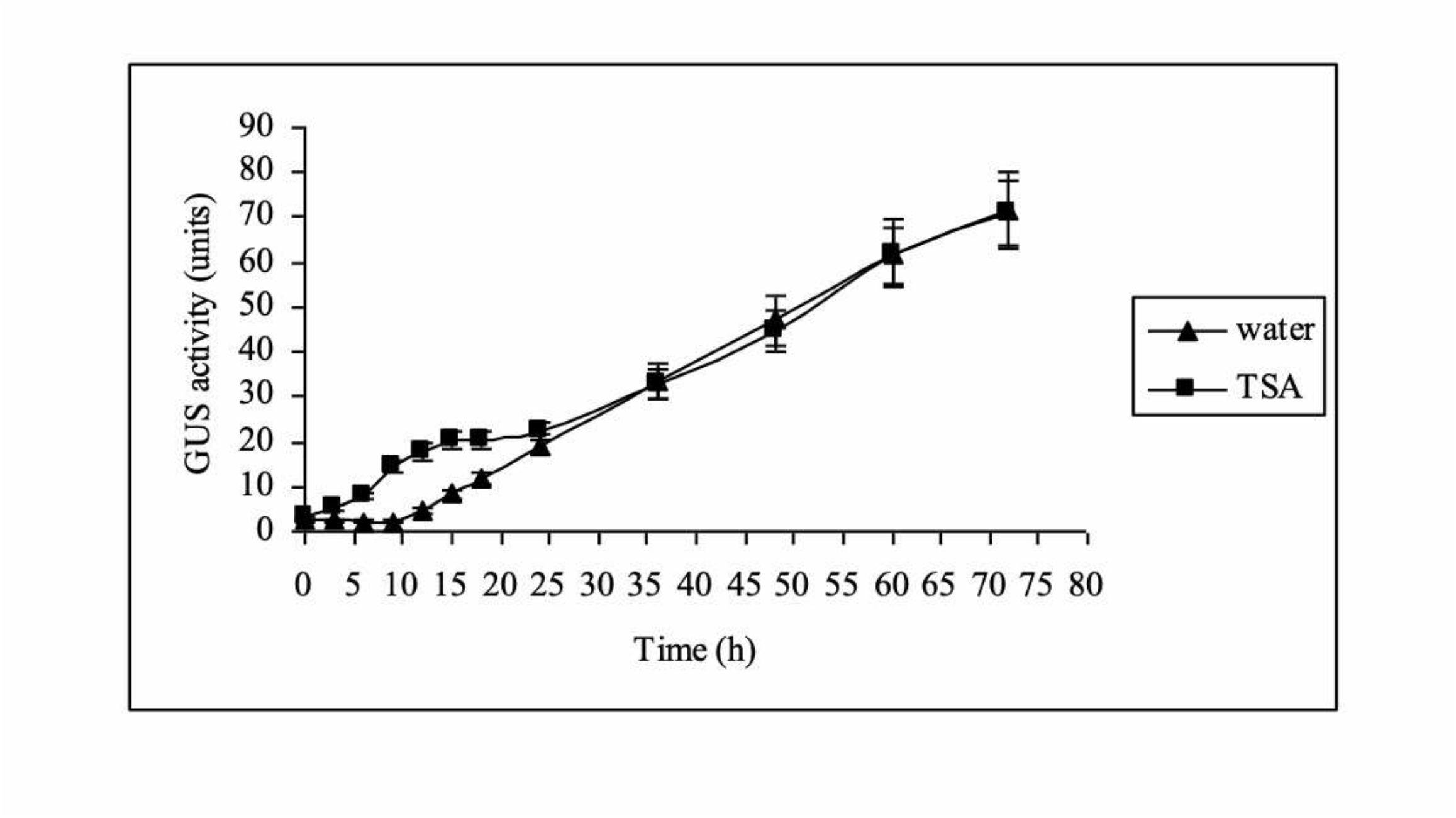
Effect of TSA on tobacco *PR-1a* expression, following induction with SA. Leaf discs from the *PR-1a*:GUS transgenic tobacco were placed in 2 mM SA for 4h only then were shifted to water or TSA for different time intervals, as indicated. The GUS assay was performed after 24 h of time completion. The kinetics of GUS expression *PR-1a* in the presence of water (▲) and TSA (■) is shown in Fig.

### Histone methylation plays a dual role in the transcriptional regulation of *PR-1a*

The role of histone methylation of nucleosome over the core promoter in *PR-1a* expression was also examined, using ChIP-qRT-PCR with antibodies specific to mono-, di- or tri-methylated H3K4, H3K9 and H4K20. A gradual increase in H3K4 me2 (Fig. 6A), H3K4me3 (Fig. 6B) and H3K9 me3 (Fig. 6F) was observed till 9h post-SA treatment coinciding with transcription activation of *PR-1a* (Fig. 4) and removal of nucleosome from the core promoter (Fig. 3). In contrast, H3K9 me1 (Fig. 6D) and me2 (Fig. 6E) were found to be enriched in the uninduced conditions and decreased subsequently post the SA treatment. H3K4 mono-methylation increased gradually up to 9h accompanies the transcriptional activation at 9h post-SA treatment (Fig. 6A). An increased trimethylation of lysine residues of H3K9 showed a dual role of histone methylation (activation and repression) in the transcriptional regulation of *PR-1a*. The methylation state of H4K20 was studied further, mono-, di- and tri-methylation of H4K20 (Fig. 6G-I) showed significantly low signals.

**Fig. 6.**
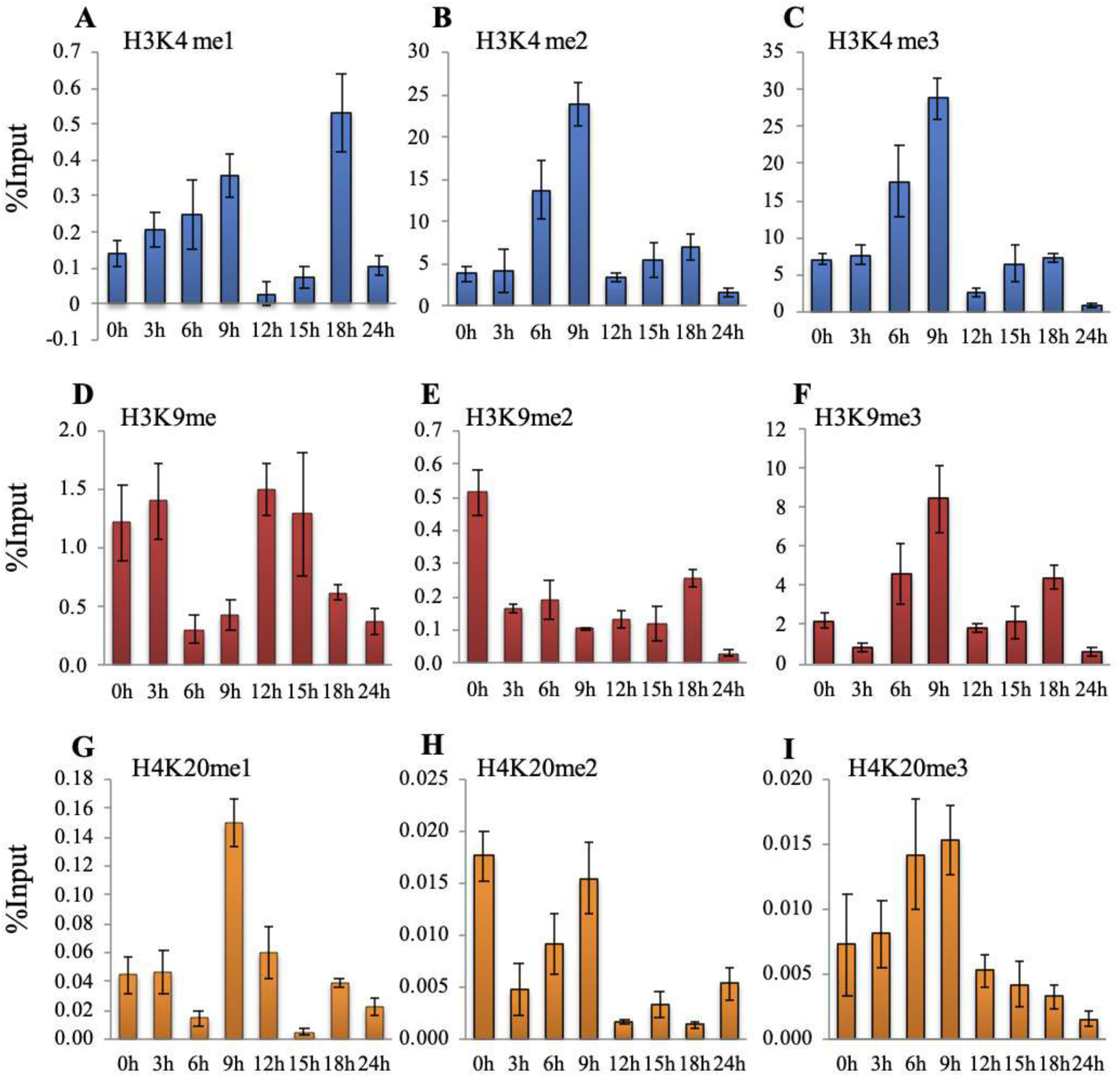
Time course analysis of the methylation status on R8 upon SA induction for different time periods. Histone methylation status on R8 was analyzed by ChIP assay using antibodies against mono-, di- and tri-methyl H3-K4, H3-K9 and H4-K20 (A-I). ChIP assay was performed using these antibodies on tobacco leaves treated with water (uninduced) or SA (induced) up to 24 h. The immunoprecipitated DNA was analyzed by qRT-PCR. The histogram represents the % input (Y-axis) at different time points (X-axis) with SD.

### The Human Lysine Specific Demethylase 1 (LSD1) like gene maintains silent state of *PR-1a* in uninduced state

To examine whether *PR1* locus genetically interacted with LSD1 like genes, we performed experiments in *Arabidopsis thaliana* (Ws) because LSD1 like mutants of tobacco plant were not available. Four putative homologs (1 to 4) of LSD1 have been reported in *A. thaliana* viz. At3G13682, At3G10390, At1G62830 and At4G16310 (Chang and Pikaard, 2005). We carried out quantitative real-time PCR of *AtPR1* transcript in these *lsd1* like mutants in the uninduced state. In all the four mutants, a high level of *AtPR1* was noticed in the uninduced state in contrast to a very low level of uninduced *AtPR1* in wild type (Fig. 7A). The mutations in LSD1 like genes (At3G10390, At1G62830, and At4G16310) led to nearly constitutive expression of *AtPR1*. The results established that the lysine-specific demethylase family was involved in giving repressed chromatin conformation to the *AtPR1* region in *A. thaliana* in the uninduced state. The results on *lsd1* mutants encouraged us to determine the recruitment of LSD1 on the core promoter region of *PR-1a*.

**Fig. 7.**
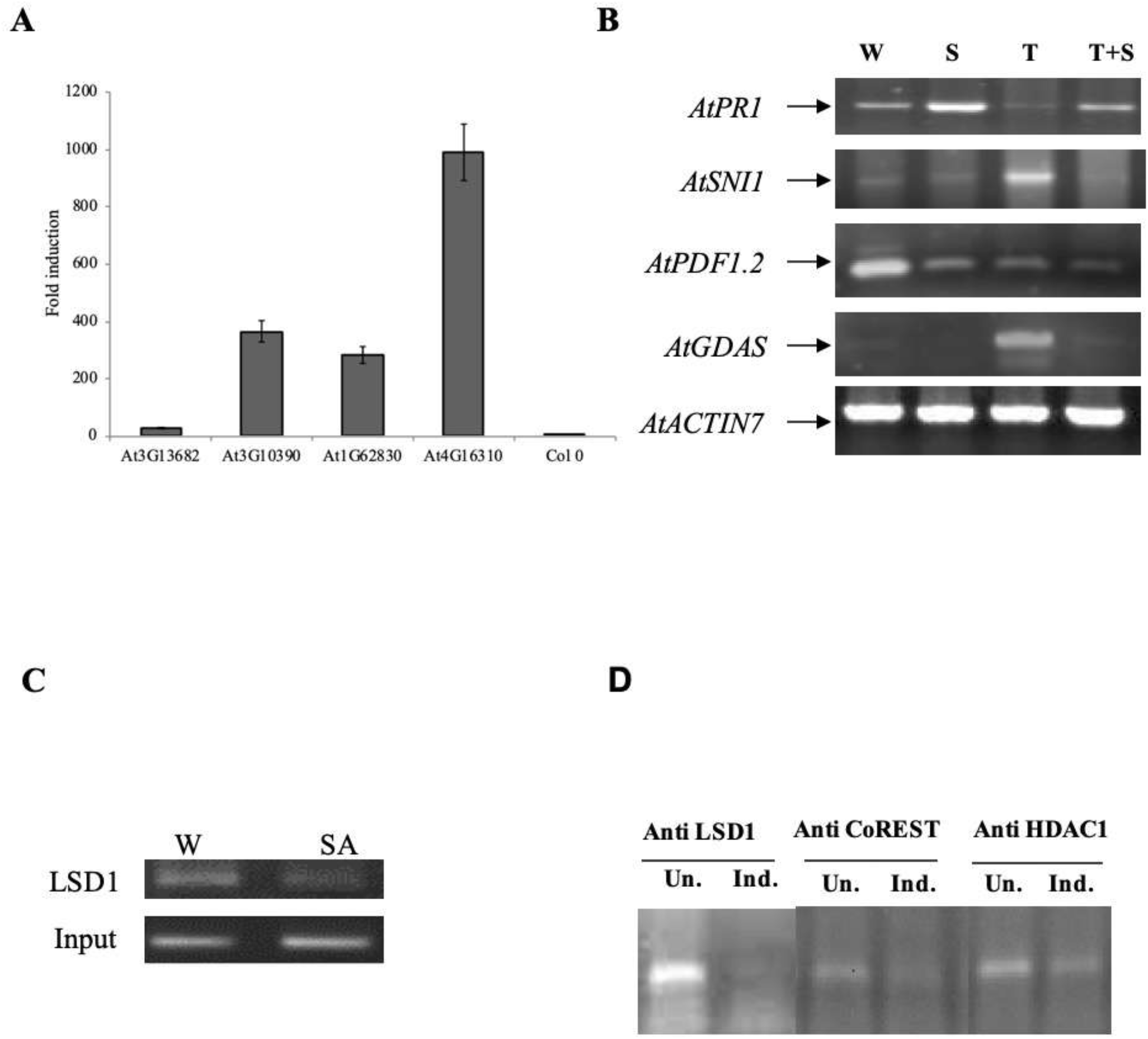
Expression of AtPR1 in mutant plants of Arabidopsis and ChIP-PCR analysis using anti-LSD1, anti-CoREST and anti-HDAC1 antibodies at *PR-1a* promoter locus. Constitutive expression of *AtPR1* in gene mutant lines of *A. thaliana* in comparison to wild type in uninduced state. (**A**). *AtPR1* expression in four LSD1 like gene mutant lines was quantified by qRT-PCR. (**B**). Expression of *AtPR1*, *AtSNI1*, *AtPDF1.2* and *AtGDAS* transcripts in *Arabidopsis*. Transcript levels were estimated by RT-PCR, 24 h after floating the leaves on water, SA, TSA and SA +TSA. The *AtACTIN7* was used as an internal control. **(C).** Presence of LSD1 on chromatin of core promoter region of *PR-1a* was analysed by ChIP assay using anti-LSD1 antibody. The immunoprecipitated DNA was analyzed by standard PCR. Input DNA was used as ChIP control. **(D).** Detection of LSD1-like complex at core promoter region of *PR-1a* in uninduced state by ChIP PCR. ChIP assay was performed by using antibodies against LSD1, CoREST and HDAC1. The representative PCR products indicate the presence of LSD1, CoREST and HDAC1, in uninduced state. Input DNA was used as ChIP control.

### TSA enhances the expression of *AtSNI1*, a negative regulator of *AtPR1*

Tobacco *PR-1a* promoter was not induced in the presence of TSA alone. To address why TSA prevents the induction of *PR-1a*, we carried out experiments on *Arabidopsis thaliana* (Columbia ecotype) because the regulators of *PR-1a* gene in tobacco have not been identified. In *A. thaliana*, a negative regulator gene of *AtPR1*, called *AtSNI1* has earlier been reported (Mosher et al., 2006). The expression of the regulatory genes was examined after treatment with SA and TSA. The *AtSNI1* gene was not activated by SA treatment but was induced by TSA (Fig. 7B). The jasmonic acid (JA) inducible *AtPDF1.2* gene was repressed by SA (Spoel et al., 2003), while TSA did not affect its expression. The TSA inducible *AtGDAS* was used as a positive control and *AtACTIN7* as an internal control in the experiments.

### LSD1-CoREST-HDAC1 complex maintains the silent state of *PR-1a*

Our results suggest the LSD1 maintains the silent state of *PR-1a* in the uninduced condition. In other studies, LSD1 was reported to be a part of the LSD1-CoREST-HDAC1 suppressor complex of neuronal genes in non-neural cells (Ballas et al., 2001). We examined whether this repressor complex was involved in maintaining the silent state of *PR-1a* also in the uninduced state. First, we checked the presence of LSD1 like protein on the core promoter region in uninduced state. ChIP analysis of *PR-1a* locus was carried out in uninduced and induced states using custom made (Supplementary information 5) anti-LSD1 specific antibody. ChIP qRT-PCR result suggested that LSD1 like protein was indeed present on the core promoter region in the uninduced state (Fig. 8A). Next, we looked for the LSD1-CoREST-HDAC1 Complex on *PR1-a* locus. We performed ChIP using again anti-LSD1 like, anti-CoREST and anti-HDAC1 specific antibodies. The results indicate the presence of CoREST and HDAC1 in the uninduced state of *PR-1a* chromatin similar as noticed for LSD1. The CoREST and HDAC1 were reduced when *PR-1a* was activated by SA (Fig. 8B).

**Fig. 8.**
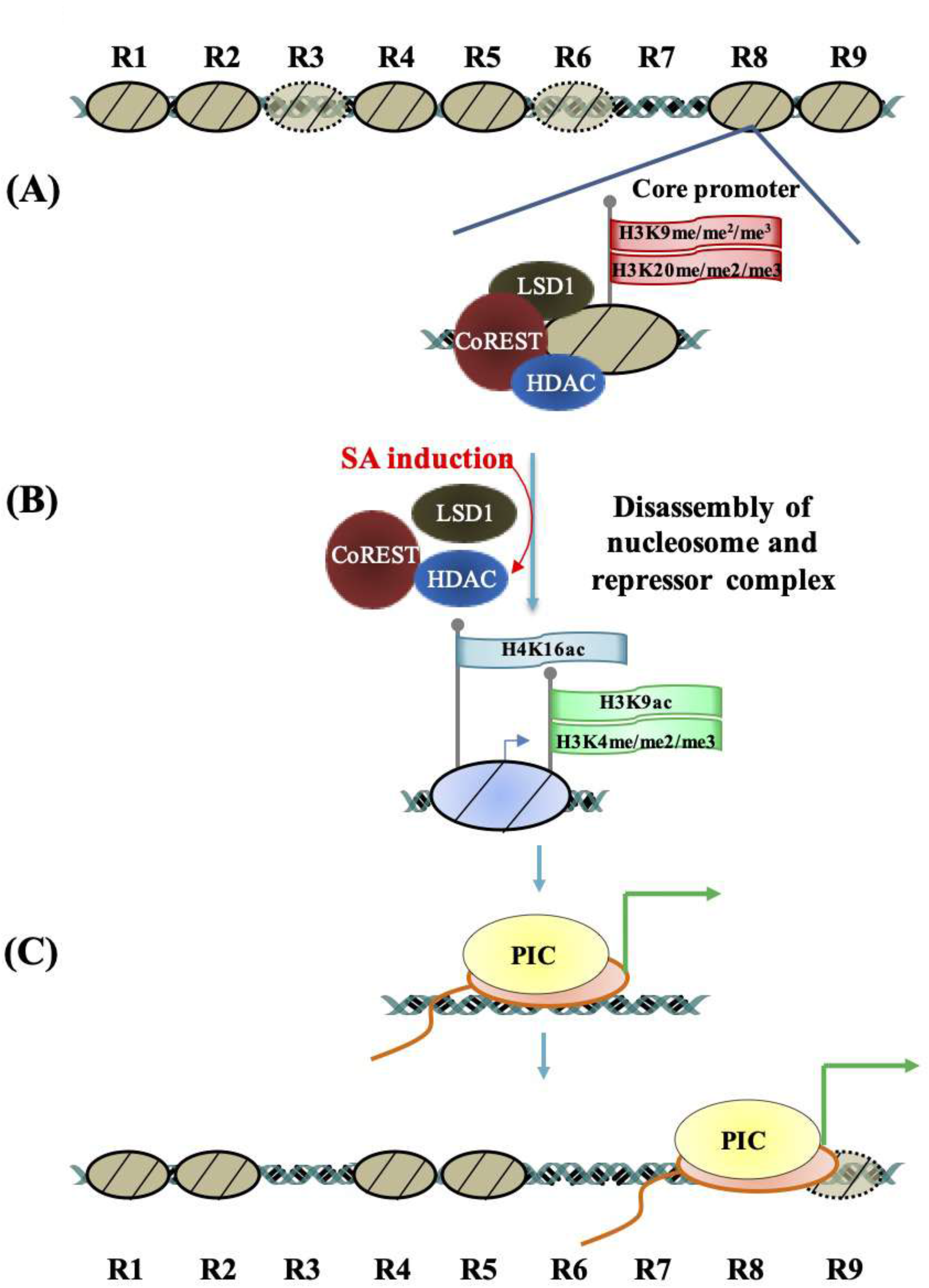
Probable Model suggesting the sequential events and ordered modifications of chromatin over the *PR-1a* promoter in tobacco leaf. Histone modifications associated with various *PR-1a* promoter states are shown. The promoter region has six distinct nucleosomes including downstream nucleosome in the repressed state, as shown in (A). The nucleosome over core promoter has repressive histone marks (mono, di and trimethylated H4-K20 and H3-K9) and LSD1-CoREST-HDAC1 repressor complex (A). Following SA mediated activation (B) of *PR-1a* promoter, the repressor complex is dissociated from the core promoter region, possibly through the recruitment of histone acetyltransferase, resulting in H3K9ac and H4K16ac. Active histone methylation marks (mono, di and trimethylated H3-K4) also increase. Acetylation at H3-K9/14 and H4-K12 lead to decrease in histone–DNA interactions eventually nucleosome disappears from the core promoter (C) region, leading to the recruitment of pre-initiation complex (PIC). The new incorporated histone codes (mono, di and trimethylation of H3-K4, and acetylation of H3-K9/14 and H4-K12) make actively transcribed *PR-1a* chromatin.

## DISCUSSION

Salicylic acid (SA) is the key signal molecule for the establishment of systemic acquired resistance (SAR). Transcripts of tobacco *PR-1a* or *AtPR1* is accumulated in response to SA signalling, which is a marker for the establishment of SAR (Loake and Grant, 2007; Vlot et al., 2009). Several efforts were made to elucidate the molecular mechanism of transcriptional regulation of *PR* genes (Kesarwani et al., 2007; Wang et al., 2009). In our present study, we focused on the epigenetic regulation core promoter nucleosome of the *PR-1a* gene and identified five nucleosomes over the promoter region of *PR-1a* in the uninduced condition spanning from the TATA-box and transcription initiation site to upstream region (as-1 like element) (Fig. 1A and B; Fig. 1S B). The nucleosome over the TATA-box is responsible for the silent state of *PR-1a* transcription in the uninduced condition (Lodhi et al., 2008) and the unmasking of the TATA-box region is crucial to establish the pre-initiation complex and recruitment of RNA polymerase II (Cairns, 2009; Juven-Gershon and Kadonaga, 2009; Kiran et al., 2006). The mechanism involving masking of the TATA-box by the nucleosome and suppression of transcription has been reported in several eukaryotic promoters (Lebel et al., 1998; Srivastava et al., 2014; Workman and Kingston, 1998). The nucleosome over the region 8 (R8) (Fig. 1S B) disappears post-SA treatment (Fig. 3; group 3 (R8)) and coincides with the *PR-1a* transcription (Fig. 4). The disappearance of the nucleosome over the R8 could be either because of nucleosome sliding (Lomvardas and Thanos, 2001) or nucleosome disassembly (Adkins et al., 2007; Boeger et al., 2004) both the mechanisms have been demonstrated in detail in different eukaryotic promoters (Boeger et al., 2005). Our native ChIP experiment using anti-H3 antibody (Fig. 2 A and B) establishes that the disappearance of the nucleosome over the R8 (group 3) could not be possible because of sliding since the region immediately downstream of the core promoter (group 4) was occupied by a nucleosome and region immediately upstream (group 2) is always free of the nucleosome. It further confirms the lack of core histone from the R8 (group 3) in the SA induced condition (Fig. 3). Thus, our results strongly support that the disappearance of the nucleosome over the R8 post-SA treatment is due to complete nucleosome disassembly. The *Anti-Silencing Function1* gene (*Asf1*) is reported to disassemble the nucleosome in budding yeast (Adkins et al., 2007). Homologs of *Asf1* have been reported from *A. thaliana* as well, suggested the possibility that nucleosome over the core region of tobacco *PR-1a* is disassembled by homologues of such genes.

Following SA induction of *PR-1a*, acetylation of H3K9/14 increased 9 h post-SA treatment (Fig. 2 A and B), similar to transgenic plants with *PR-1a:GUS*, the expression of GUS protein was detected at 10 h post SA induction (Fig. 5) (Lodhi et al., 2008). A rapid transient increase in acetylation of H3K9/14 at 9 h and a slight increase in tri-methylation of H3K4 in the activation of *PR-1a* transcription at the same time (Fig. 4) indicate that the H3-K9/K14 acetylation is required for the active state of *PR-1a* core promoter. The acetylation of H3-K9/14 has been reported in the activation of *RBCS*-1A and *IAA3* genes (Benhamed et al., 2006). Microarray analyses in tobacco and *A. thaliana* seedlings show that TSA induces changes in gene expression and affects histone acetylation in specific genes (Chang and Pikaard, 2005; Chua et al., 2004). In *A. thaliana*, histone deacetylase *AtHD1* (also called HDA19) is involved in the regulation of pathogen response genes (Zhou et al., 2005). We observed TSA mediated suppression of *AtPR1* transcription (Fig. 7B) and also inhibition in nucleosome modeling at the core promoter (Fig. 3) when TSA was provided along with SA. These results were surprising in the context of the importance of H3K9/14 and H4K16 acetylation required for *PR-1a* activation (Fig. 2A-B). One possible explanation could be that TSA mediated suppression of *PR-1a* is indirect by higher expression of a negative regulator of *PR-1a* locus.

Modification of the histone H3K4 di- and tri-methylation also enrich till 9h post-induction and positively correlate transcriptional activation (Fig. 6B-C). Mono methylation of H3K4 is initially very little enrichment and its transient mild enrichment till 9h at the *PR-1a* promoter. Earlier reports also suggest that the presence of H3K4me2 and H3K4me3 in plants is usually correlated with the active transcription of the highly expressed genes, whereas H3K4me1 is distributed within transcribed regions (Zhang et al., 2009). Our results also suggested that histone modification such as mono and dimethylation at lysine 9 and 20 of H3 and H4 respectively were found increased in the uninduced state of *PR-1a* (Fig. 6D, E, and H). This agrees with the earlier reports that H4K20 methylation results in the repression of genes, which is associated with silent chromatin and inhibits acetylation of H4K16 (Karachentsev et al., 2005; Sarg et al., 2004; Sims et al., 2003). Following SA induction, a decrease in H3K9 mono- and di-methylation suggested their involvement in repressing the locus in the uninduced state, also reported by several other studies (Bernatavichute et al., 2008; Fuchs et al., 2006; Jackson et al., 2004; Johnson et al., 2004; Lippman et al., 2004; Mathieu et al., 2005). This decrease may be their conversion to the trimethylated state as shown by the H3K9 trimethylation enrichment, which is a mark for transcriptional activation (Turck et al., 2007). Lack of H4K20 methylation in transcriptionally active regions has also been reported in the Drosophila male X chromosome as the methylation of H4K20 precludes acetylation of the neighboring H4K16, both processes being competitive (Nishioka et al., 2002). However, ORC1-dependent gene activation in plants is associated with an increase in H4 acetylation and H4K20 trimethylation (de la Paz Sanchez and Gutierrez, 2009). Moreover, monomethylated H4K20 is associated with heterochromatin, and di- and tri-methylated H4K20 are associated with euchromatin in Arabidopsis (Naumann et al., 2005).

It is conceivable that the loss in di- and tri-methylation of H4K20 and di-methylation of H3K4 in *PR-1a* in the induced state results from enzymatic demethylation. A human LSD1 that demethylates mono and di-methylated H3K4 has been identified (Chang and Pikaard, 2005), suggesting involvement of LSD1 like genes in tobacco for demethylation of the di-methylated H3K4. Full enzymatic activity of LSD1 requires its association with other proteins, such as CoREST (restin corepressor) complex, indicating that regulatory subunits can have a role in modulating demethylase activity (Chang and Pikaard, 2005; Lee et al., 2005). The presence of a nucleosome over the core promoter has often been associated with transcriptional silencing of genes (Lebel et al., 1998; Srivastava et al., 2014). Our study shows that five nucleosomes cover the promoter region of *PR-1a* including a nucleosome over the downstream region (core promoter) or upstream activator region (covers *as-1-*like element responsible for induction) (Butterbrodt et al., 2006). After induction, the nucleosome over the core promoter disassembles and provides the access to transcription initiation machinery on nucleosome-free core promoter region. In conclusion, we suggest nucleosome association with LSD1-CoREST-HDAC1 suppressor like complex maintain the silent state of *PR-1a* locus (Fig. 8).

## Supporting information

List of primers used in this study

Position of nucleosomes on promoter of PR-1a in uninduced state

## ACKNOWLEDGMENTS

The authors are grateful to Council of Scientific and Industrial Research (CSIR) for Junior/Senior Research Fellowship to Niraj Lodhi and Department of Sciences and Technology, Government of India for the research grant, J.C. Bose and Research Fellowships to Dr. Rakesh Tuli.

## SUPPLEMENTARY INFORMATION

**Table S1** List of primers used in this study.

**Figure S1** Position of nucleosomes on promoter of PR-1a in uninduced state.

## REFERENCES

Adkins, M.W., Williams, S.K., Linger, J., Tyler, J.K. (2007) Chromatin disassembly from the PHO5 promoter is essential for the recruitment of the general transcription machinery and coactivators. Molecular and Cellular Biology, 27(18), 6372–6382.

Ahmad, K. and Henikoff, S. (2002) The histone variant H3.3 marks active chromatin by replication-independent nucleosome assembly. Molecular Cell, 9(6), 1191–1200.

Ballas, N., Battaglioli, E., Atouf, F., Andres, M.E., Chenoweth, J., Anderson, M.E., Burger, C., Moniwa, M., Davie, J.R., Bowers, W.J., Federoff, H.J., Rose, D.W., Rosenfeld, M.G., Brehm, P., Mandel, G. (2001) Regulation of neuronal traits by a novel transcriptional complex. Neuron, 31(3), 353–365.

Bastow, R., Mylne, J.S., Lister, C., Lippman, Z., Martienssen, R.A., Dean, C. (2004) Vernalization requires epigenetic silencing of FLC by histone methylation. Nature, 427(6970), 164–167.

Benhamed, M., Bertrand, C., Servet, C., Zhou, D.X. (2006) Arabidopsis GCN5, HD1, and TAF1/HAF2 interact to regulate histone acetylation required for light-responsive gene expression. The Plant Cell, 18(11), 2893–2903.

Bernatavichute, Y.V., Zhang, X., Cokus, S., Pellegrini, M., Jacobsen, S.E. (2008) Genome-wide association of histone H3 lysine nine methylation with CHG DNA methylation in Arabidopsis thaliana. PLoS One, 3(9), e3156.

Boeger, H., Bushnell, D.A., Davis, R., Griesenbeck, J., Lorch, Y., Strattan, J.S., Westover, K.D., Kornberg, R.D. (2005) Structural basis of eukaryotic gene transcription. FEBS Letters, 579(4), 899–903.

Boeger, H., Griesenbeck, J., Strattan, J.S., Kornberg, R.D. (2004) Removal of promoter nucleosomes by disassembly rather than sliding in vivo. Molecular Cell, 14(5), 667–673.

Brand, M., Rampalli, S., Chaturvedi, C.P., Dilworth, F.J. (2008) Analysis of epigenetic modifications of chromatin at specific gene loci by native chromatin immunoprecipitation of nucleosomes isolated using hydroxyapatite chromatography. Nature Protocols, 3(3), 398–409.

Butterbrodt, T., Thurow, C., Gatz, C. (2006) Chromatin immunoprecipitation analysis of the tobacco PR-1a- and the truncated CaMV 35S promoter reveals differences in salicylic acid-dependent TGA factor binding and histone acetylation. Plant Mol Biol, 61(4-5), 665–674.

Cairns, B.R. (2009) The logic of chromatin architecture and remodelling at promoters. Nature, 461(7261), 193–198.

Chang, S. and Pikaard, C.S. (2005) Transcript profiling in Arabidopsis reveals complex responses to global inhibition of DNA methylation and histone deacetylation. Journal of Biological Chemistry, 280(1), 796–804.

Chua, Y.L., Mott, E., Brown, A.P., MacLean, D., Gray, J.C. (2004) Microarray analysis of chromatin-immunoprecipitated DNA identifies specific regions of tobacco genes associated with acetylated histones. The Plant Journal, 37(6), 789–800.

Chua, Y.L., Watson, L.A., Gray, J.C. (2003) The transcriptional enhancer of the pea plastocyanin gene associates with the nuclear matrix and regulates gene expression through histone acetylation. The Plant Cell, 15(6), 1468–1479.

de la Paz Sanchez, M. and Gutierrez, C. (2009) Arabidopsis ORC1 is a PHD-containing H3K4me3 effector that regulates transcription. Proceedings of the National Academy of Sciences, 106(6), 2065–2070.

Fischle, W., Wang, Y., Allis, C.D. (2003) Histone and chromatin cross-talk. Current Opinion in Cell Biology, 15(2), 172–183.

Fuchs, J., Demidov, D., Houben, A., Schubert, I. (2006) Chromosomal histone modification patterns--from conservation to diversity. Trends in Plant Science, 11(4), 199–208.

Jackson, J.P., Johnson, L., Jasencakova, Z., Zhang, X., PerezBurgos, L., Singh, P.B., Cheng, X., Schubert, I., Jenuwein, T., Jacobsen, S.E. (2004) Dimethylation of histone H3 lysine 9 is a critical mark for DNA methylation and gene silencing in Arabidopsis thaliana. Chromosoma, 112(6), 308–315.

Jenuwein, T. and Allis, C.D. (2001) Translating the histone code. Science, 293(5532), 1074–1080.

Johnson, L., Mollah, S., Garcia, B.A., Muratore, T.L., Shabanowitz, J., Hunt, D.F., Jacobsen, S.E. (2004) Mass spectrometry analysis of Arabidopsis histone H3 reveals distinct combinations of post-translational modifications. Nucleic Acids Res, 32(22), 6511–6518.

Juven-Gershon, T. and Kadonaga, J.T. (2009) Regulation of gene expression via the core promoter and the basal transcriptional machinery. Developmental Biology, 339(2), 225–229.

Karachentsev, D., Sarma, K., Reinberg, D., Steward, R. (2005) PR-Set7-dependent methylation of histone H4 Lys 20 functions in repression of gene expression and is essential for mitosis. Genes & Development, 19(4), 431–435.

Kesarwani, M., Yoo, J., Dong, X. (2007) Genetic interactions of TGA transcription factors in the regulation of pathogenesis-related genes and disease resistance in Arabidopsis. Plant Physiology, 144(1), 336–346.

Kiran, K., Ansari, S.A., Srivastava, R., Lodhi, N., Chaturvedi, C.P., Sawant, S.V., Tuli, R. (2006) The TATA-box sequence in the basal promoter contributes to determining light-dependent gene expression in plants. Plant Physiology, 142(1), 364–376.

Lebel, E., Heifetz, P., Thorne, L., Uknes, S., Ryals, J., Ward, E. (1998) Functional analysis of regulatory sequences controlling PR-1 gene expression in Arabidopsis. The Plant Journal, 16(2), 223–233.

Lee, M.G., Wynder, C., Cooch, N., Shiekhattar, R. (2005) An essential role for CoREST in nucleosomal histone 3 lysine 4 demethylation. Nature, 437(7057), 432–435.

Lippman, Z., Gendrel, A.V., Black, M., Vaughn, M.W., Dedhia, N., McCombie, W.R., Lavine, K., Mittal, V., May, B., Kasschau, K.D., Carrington, J.C., Doerge, R.W., Colot, V., Martienssen, R. (2004) Role of transposable elements in heterochromatin and epigenetic control. Nature, 430(6998), 471–476.

Loake, G. and Grant, M. (2007) Salicylic acid in plant defence--the players and protagonists. Current Opinion in Plant Biology, 10(5), 466–472.

Lodhi, N., Ranjan, A., Singh, M., Srivastava, R., Singh, S.P., Chaturvedi, C.P., Ansari, S.A., Sawant, S.V., Tuli, R. (2008) Interactions between upstream and core promoter sequences determine gene expression and nucleosome positioning in tobacco PR-1a promoter. Biochimica et Biophysica Acta (BBA) - Gene Regulatory Mechanisms, 1779(10), 634–644.

Lomvardas, S. and Thanos, D. (2001) Nucleosome sliding via TBP DNA binding in vivo. Cell, 106(6), 685–696.

Lomvardas, S. and Thanos, D. (2002) Modifying gene expression programs by altering core promoter chromatin architecture. Cell, 110(2), 261–271.

Mathieu, O., Probst, A.V., Paszkowski, J. (2005) Distinct regulation of histone H3 methylation at lysines 27 and 9 by CpG methylation in Arabidopsis. The EMBO Journal, 24(15), 2783–2791.

Metzger, E., Wissmann, M., Yin, N., Muller, J.M., Schneider, R., Peters, A.H., Gunther, T., Buettner, R., Schule, R. (2005) LSD1 demethylates repressive histone marks to promote androgen-receptor-dependent transcription. Nature, 437(7057), 436–439.

Mosher, R.A., Durrant, W.E., Wang, D., Song, J., Dong, X. (2006) A comprehensive structure-function analysis of Arabidopsis SNI1 defines essential regions and transcriptional repressor activity. The Plant Cell, 18(7), 1750–1765.

Naumann, K., Fischer, A., Hofmann, I., Krauss, V., Phalke, S., Irmler, K., Hause, G., Aurich, A.C., Dorn, R., Jenuwein, T., Reuter, G. (2005) Pivotal role of AtSUVH2 in heterochromatic histone methylation and gene silencing in Arabidopsis. The EMBO Journal, 24(7), 1418–1429.

Nishioka, K., Rice, J.C., Sarma, K., Erdjument-Bromage, H., Werner, J., Wang, Y., Chuikov, S., Valenzuela, P., Tempst, P., Steward, R., Lis, J.T., Allis, C.D., Reinberg, D. (2002) PR-Set7 is a nucleosome-specific methyltransferase that modifies lysine 20 of histone H4 and is associated with silent chromatin. Molecular Cell, 9(6), 1201–1213.

Santos-Rosa, H. and Caldas, C. (2005) Chromatin modifier enzymes, the histone code and cancer. European Journal of Cancer, 41(16), 2381–2402.

Sarg, B., Helliger, W., Talasz, H., Koutzamani, E., Lindner, H.H. (2004) Histone H4 hyperacetylation precludes histone H4 lysine 20 trimethylation. Journal of Biological Chemistry, 279(51), 53458–53464.

Shahbazian, M.D. and Grunstein, M. (2007) Functions of site-specific histone acetylation and deacetylation. Annual Review of Biochemistry, 76, 75–100.

Shogren-Knaak, M. and Peterson, C.L. (2006) Switching on chromatin: mechanistic role of histone H4-K16 acetylation. Cell Cycle, 5(13), 1361–1365.

Sims, R.J., 3rd, Nishioka, K., Reinberg, D. (2003) Histone lysine methylation: a signature for chromatin function. Trends in Genetics, 19(11), 629–639.

Spoel, S.H., Koornneef, A., Claessens, S.M., Korzelius, J.P., Van Pelt, J.A., Mueller, M.J., Buchala, A.J., Métraux, J.P., Brown, R., Kazan, K., Van Loon, L.C., Dong, X., Pieterse, C.M. (2003) NPR1 modulates cross-talk between salicylate- and jasmonate-dependent defense pathways through a novel function in the cytosol. The Plant Cell, 15(3), 760–770.

Srivastava, R. and Ahn, S.H. (2015) Modifications of RNA polymerase II CTD: Connections to the histone code and cellular function. Biotechnology Advances, 33(6, Part 1), 856–872.

Srivastava, R., Rai, K.M., Srivastava, M., Kumar, V., Pandey, B., Singh, S.P., Bag, S.K., Singh, B.D., Tuli, R., Sawant, S.V. (2014) Distinct Role of Core Promoter Architecture in Regulation of Light-Mediated Responses in Plant Genes. Molecular Plant, 7(4), 626–641.

Srivastava, R., Singh, U.M., Dubey, N.K. (2016) Histone Modifications by different histone modifiers: insights into histone writers and erasers during chromatin modification. Journal of Biological Sciences and Medicine, 2(1), 45–54.

Struhl, K. (1998) Histone acetylation and transcriptional regulatory mechanisms. Genes & Development, 12(5), 599–606.

Sung, S. and Amasino, R.M. (2004) Vernalization and epigenetics: how plants remember winter. Current Opinion in Plant Biology, 7(1), 4–10.

Tsuji, H., Saika, H., Tsutsumi, N., Hirai, A., Nakazono, M. (2006) Dynamic and reversible changes in histone H3-Lys4 methylation and H3 acetylation occurring at submergence-inducible genes in rice. Plant and Cell Physiology, 47(7), 995–1003.

Tsukada, Y., Fang, J., Erdjument-Bromage, H., Warren, M.E., Borchers, C.H., Tempst, P., Zhang, Y. (2006) Histone demethylation by a family of JmjC domain-containing proteins. Nature, 439(7078), 811–816.

Turck, F., Roudier, F., Farrona, S., Martin-Magniette, M.L., Guillaume, E., Buisine, N., Gagnot, S., Martienssen, R.A., Coupland, G., Colot, V. (2007) Arabidopsis TFL2/LHP1 specifically associates with genes marked by trimethylation of histone H3 lysine 27. PLOS Genetics, 3(6), e86.

Turner, B.M. (2000) Histone acetylation and an epigenetic code. Bioessays, 22(9), 836–845.

Vlot, A.C., Dempsey, D.A., Klessig, D.F. (2009) Salicylic Acid, a multifaceted hormone to combat disease. Annual Review of Phytopathology, 47, 177–206.

Wang, X., Basnayake, B.M., Zhang, H., Li, G., Li, W., Virk, N., Mengiste, T., Song, F. (2009) The Arabidopsis ATAF1, a NAC transcription factor, is a negative regulator of defense responses against necrotrophic fungal and bacterial pathogens. Molecular Plant-Microbe Interactions, 22(10), 1227–1238.

Whetstine, J.R., Nottke, A., Lan, F., Huarte, M., Smolikov, S., Chen, Z., Spooner, E., Li, E., Zhang, G., Colaiacovo, M., Shi, Y. (2006) Reversal of histone lysine trimethylation by the JMJD2 family of histone demethylases. Cell, 125(3), 467–481.

Workman, J.L. and Kingston, R.E. (1998) Alteration of nucleosome structure as a mechanism of transcriptional regulation. Annual Review of Biochemistry, 67, 545–579.

Yamane, K., Toumazou, C., Tsukada, Y., Erdjument-Bromage, H., Tempst, P., Wong, J., Zhang, Y. (2006) JHDM2A, a JmjC-containing H3K9 demethylase, facilitates transcription activation by androgen receptor. Cell, 125(3), 483–495.

Zhang, X., Bernatavichute, Y.V., Cokus, S., Pellegrini, M., Jacobsen, S.E. (2009) Genome-wide analysis of mono-, di- and trimethylation of histone H3 lysine 4 in Arabidopsis thaliana. Genome Biology, 10(6), R62.

Zhou, C., Zhang, L., Duan, J., Miki, B., Wu, K. (2005) HISTONE DEACETYLASE19 is involved in jasmonic acid and ethylene signaling of pathogen response in Arabidopsis. The Plant Cell, 17(4), 1196–1204.

