## Supplementary figures and images for "Epigenetic Malleability at Core Promoter Regulates Tobacco PR-1a Expression after Salicylic Acid Treatment"

### List of primers used in this study

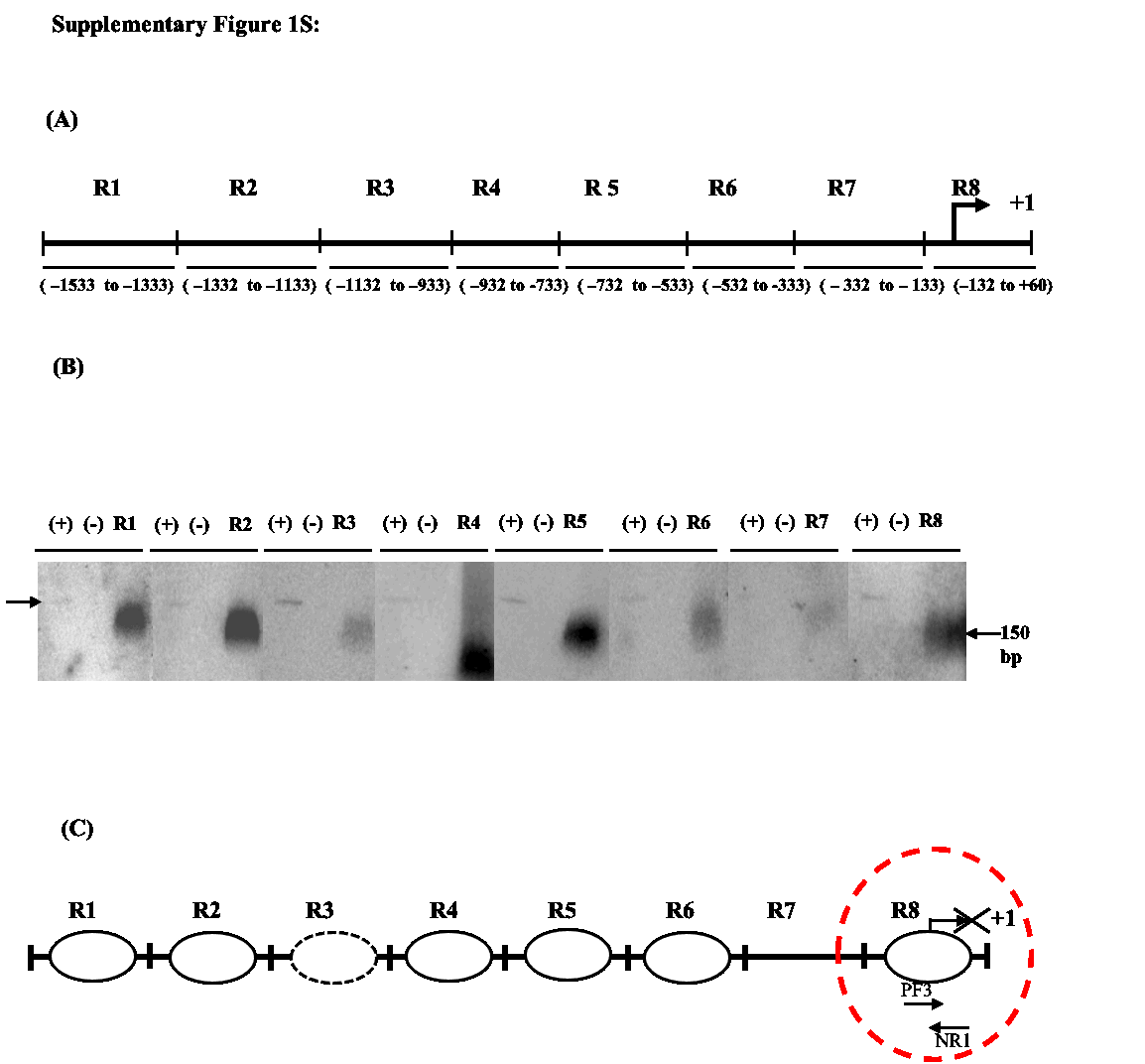
