## Supplementary material for "Epigenetic Malleability at Core Promoter Regulates Tobacco PR-1a Expression after Salicylic Acid Treatment": Position of nucleosomes on promoter of PR-1a in uninduced state

**For Cloning of *PR-1a* promoter**

1. PRF*	5’ TCA ACT **CTG CAG** GTC GAC GGA CTA AGA TTA CGA GGA T 3’

2. PRR**	5’ TAC TAG **TCT AGA** ATC GAT GAC TAT AGG AGA 3’

**For detection of gene expression**

3. ATPRF	5’ CGA ACA CGT GCA ATG GAG TTT 3’

4. ATPRR	5’ CCC ACG AGG ATC ATA GTT GCA A 3’

5. AT*ACTIN*7F	5’ AAG TCA TAA CCA TCG GAG CTG 3’

6. AT*ACTIN*7R	5’ ACC AGA TAA GAC AAG ACA CAC 3’

7. NPRF	5’ATG GGA TTT GTT CTC TTT TCA C 3’

8. NPRR	5’TAG TAT GGA CTT TCG CCT CT 3’

9. UBIF	5’ GAA GCA GCT CGA GGA TGG AA 3’

10.UBIR	5’ CCA CGG AGA CGG AGG ACA A 3’

**For detection of nucleosome**

11. PF3	5’ TGA TAT TAC CAT GTC AAA AAA TTT AGT 3’

12. NR1	5’ CAA TTG TGA AAA GAG AAC AAA T 3’

13. NPAF1	5’ TAT AAA TAT GGA AGT AAA AAT TAA TC 3’

14. NPAR1	5’ TAA TTT TAA ATT GTC AAT GCA TGA 3’

15. NPAF5	5’ CTG ATA GAT CAA AAA AGT GTT TAA CT 3’

16. NPAR5	5’ ATG TAA TAT ATC CTG TTA TAG ATA A 3’

17. AGF	5’ GCC GTT TCA AAT TAA GTC TAT TCT G 3’

18. AGR	5’ TCG TAG GAG GAA TTA GTT AGT GGT G 3’

* PstI

**XbaI

Bold nucleotides indicate restriction sites

**Supplementary Table 1**

**Primers used in this study:**
